# Mapping Cellular Response to Destabilized Transthyretin Reveals Cell- and Amyloidogenic Protein-Specific Signatures

**DOI:** 10.1101/2022.08.17.504308

**Authors:** Sabrina Ghosh, Carlos Villacorta-Martin, Jonathan Lindstrom-Vautrin, Devin Kenney, Carly S. Golden, Camille V. Edwards, Vaishali Sanchorawala, Lawreen H. Connors, Richard M. Giadone, George J. Murphy

## Abstract

In ATTR amyloidosis, transthyretin (TTR) protein is secreted from the liver and deposited as toxic aggregates at downstream target tissues. Despite recent advancements in treatments for ATTR amyloidosis, the mechanisms underlying misfolded TTR-mediated cellular damage remain elusive. In an effort to define early events of TTR-associated stress, we exposed neuronal (SH-SY5Y) and cardiac (AC16) cells to wild-type and destabilized TTR variants (TTR^V122I^ and TTR^L55P^) and performed transcriptional (RNAseq) and epigenetic (ATACseq) profiling. We subsequently compared TTR-responsive signatures to cells exposed to destabilized antibody light chain protein associated with AL amyloidosis as well as ER stressors (thapsigargin, heat shock). In doing so, we observed overlapping, yet distinct cell type- and amyloidogenic protein-specific signatures, suggesting unique responses to each amyloidogenic variant. Moreover, we identified chromatin level changes in AC16 cells exposed to mutant TTR which resolved upon pre-incubation with kinetic stabilizer tafamidis. Collectively, these data provide insight into the mechanisms underlying amyloid-mediated cellular damage and provide a robust resource representing cellular responses to aggregation-prone proteins and ER stress.

## Introduction

The systemic amyloid diseases are a class of disorders in which effector organs produce proteins that misfold and aggregate, travel throughout circulation, and deposit extracellularly at distal target sites^1–3^. Two of the most prevalent examples of systemic amyloid diseases include transthyretin amyloidosis (ATTR amyloidosis) and light chain amyloidosis (AL amyloidosis). In ATTR amyloidosis, the liver produces amyloidogenic transthyretin (TTR), while in AL amyloidosis, malignant plasma cells of the bone marrow secrete destabilized antibody light chain (LC) with high propensity to misfold^4–6^. In both diseases, misfolding TTRs and LCs travel throughout circulation and deposit extracellularly at downstream target organs (including the heart and peripheral nerves for ATTR amyloidosis, and renal tissue for AL amyloidosis), ultimately resulting in organ dysfunction^4–6^.

Despite the recent development of therapeutics for diseases like ATTR amyloidosis^1–6^, it remains unclear how deposition of extracellular, aggregated TTRs and LCs damage target cell types. At the same time, there is large diversity in the pathology of both ATTR and AL amyloidosis, with some patients exhibiting amyloid deposition and damage at peripheral nerves, while others develop significant cardiac disease^4–6^. In the case of ATTR amyloidosis, for example, the TTR^V122I^ mutation predominantly leads to cardiac amyloid deposition and dysfunction, while TTR^L55P^ damages and disrupts the functionality of nervous tissue^7,8^. Understanding the earliest responses to extracellular exposure of destabilized TTRs and LCs could provide insight into mechanisms of protein-mediated cell death as well as the identification of putative biomarkers for these difficult-to-diagnose diseases.

Recently, ER stress and related signaling pathways have been implicated in many systemic amyloid diseases including both ATTR and AL amyloidosis at multiple sites of pathogenesis – from effector organs producing misfolding-prone proteins, to target cell types interacting with large, insoluble amyloid^9–13^. Moreover, the response of cells to ER stress has been linked to both disease and aging across species and tissues^14–16^. Two robust stressors used to stimulate ER stress and related protein homeostasis (proteostasis) pathways in vitro include heat shock (HS) and thapsigargin. Activation of the HS response (HSR) results in trimerization of transcription factor HSF1, nuclear translocation of the HSF1 complex, and activation of transcriptional networks constituting upregulation of heat shock proteins (HSPs) important in ensuring proper folding of proteins^9,17^. Thapsigargin, on the other hand, is a SERCA kinase inhibitor which results in increased accumulation of intracellular calcium, leading to global activation of the unfolded protein response (UPR). Through mechanisms similar to the HSR, UPR activation leads to nuclear translocation of several transcription factors (cleaved ATF6, spliced XBP1, and ATF4 downstream of PERK), stimulation of related transcriptional networks, and ultimately, activation of functional pathways to respond to ER stress (e.g. upregulation of chaperone genes)^14,18^.

Cells respond to a variety of stimuli including xenobiotic, oxidative, and ER stress through complex intracellular signaling pathways^14,15,19,20^. Activation of these pathways can result in reprogramming of transcriptional networks in an effort to maintain function during acute stress, disease, and aging^1,9,10^. As many stress response pathways share downstream convergent mechanisms (e.g. signaling effector molecules, transcription factors, gene signatures, etc.), identifying commonalities and differences in response to diverse insults may uncover novel therapeutic avenues for ameliorating the toxicity of diseases with similar underlying etiologies.

Here, we employ bulk RNAseq and ATACseq to profile transcriptional and chromatin level changes resulting from exposure to cardiomyopathy- and peripheral neuropathy-associated TTRs (TTR^V122I^ and TTR^L55P^, respectively), destabilized LC, thapsigargin, and HS in immortalized neuronal SH-SY5Y and cardiac AC16 cell lines. In doing so, we compare the transcriptional and chromatin level responses of each cell type to each stressor. Through these efforts, we generated comprehensive datasets cataloging SH-SY5Y and AC16 cell responses to destabilized TTR and LC proteins and robust ER stressors. Lastly, we provide evidence that cells respond to each stressor via cell type- and stress-specific signatures, including different amyloidogenic proteins as well as pathologic variants of the same proteins.

## Materials and Methods

### Quantifying protein concentration via Bradford assay

Concentration of recombinant human TTRs was determined via Bradford assay. Samples were thawed on ice and diluted 1:1000 in 1X Bradford reagent (BioRad, Hercules, CA, Cat. No. 500-0006). Standard curve was determined via serial dilutions of 1 mg/mL BSA diluted in H2O. All samples, including standards, were incubated at room temperature for ~10 minutes. Absorbance was determined at 595 nm. All samples were run in technical duplicate.

### Stress conditions for cells: heat shock, thapsigargin, and recombinant TTR and LC exposure

For heat shock condition, cells were plated at 4×10^5^ cells/mL. 24 hours later, cells were incubated at 42°C for 75 minutes, and harvested using 0.05% trypsin. For thapsigargin exposure, cells were plated at 2×10^5^ cells/mL. 24 hours later, cells were exposed to 500 nM thapsigargin (Millipore Sigma, Cat. No. T9033) for 20 hours and harvested as previously stated. For rhTTR exposure for transcriptomic profiling, cells were plated at 2×10^5^ cells/mL. 24 hours later, respective recombinant protein (0.2 mg/mL TTR, 0.02 mg/mL LC) in the absence and presence of 10 µM tafamidis were added. After incubating cells at 37°C for 48 hours, cells were harvested for RNA or gDNA extraction.

### Culturing SH-SY5Y and AC16 cells

SH-SY5Y cells (ATCC, Manassas, VA, Cat. No. CRL-2266) were maintained in 1:1 EMEM and DMEM F/12 with 10% FBS. AC16 (Millipore Sigma, Burlington, MA, Cat. No. SCC109) cells were cultured in DMEM/F12 with 12.5% FBS. All cells were maintained at 37°C and 5% CO2 and grown in media supplemented with 1X primocin (Reprocell, Boston, MA, Cat. No. 04-0012-02).

### RNA-seq profiling and differential gene expression analysis

Total RNA was extracted using a RNeasy mini extraction kit (Qiagen, Hilden, Germany, Cat. No. 74104) and submitted for RNAseq via Novogene Co. (Sacramento, CA). The quality of the raw data was assessed using FastQC v.0.11.7. The sequence reads were aligned to the human genome reference (GRCh38) using STAR v.2.6.0c21. Counts per gene were summarized using the featureCounts function from the subread package v.1.6.222. After the summarization of counts per gene using featureCounts, the samples proceeded to downstream analysis. The edgeR package v.3.25.10 was used to import, organize, and filter the counts^23^. The matrix of counts per gene per sample was initially analyzed via DESeq2 and later analyzed using the limma/voom normalization method. Genes were filtered based on the standard edgeR filtration method using the default parameters for the “filterByExpr'' function. After exploratory data analysis (Glimma v. 1.11.1), contrasts for differential expression testing were done to compare naive and experimental samples. The limma package v.3.42.224 with its voom method, namely, linear modelling and empirical Bayes moderation was used to test differential expression (moderate t-test). P-values were adjusted for multiple testing using Benjamini-Hochberg correction (false discovery rate-adjusted p-value; FDR). Differentially expressed genes for each comparison were visualized using Glimma v. 1.11.125, and FDR<0.05 was set as the threshold for determining significant differential gene expression. Functional predictions were performed using the fgsea v.1.12.0 package for gene set analysis^26^.

### ATACseq sequencing, quality control, and data analysis

For ATACseq, gDNA was isolated from cells via DNeasy Blood and Tissue Kit (Qiagen, Cat. No. 69504) and processed as described in Buenrostro et al. 2013. To analyze our ATACSeq data we used the esATAC pipeline^27^. The pipeline consists of three major functions: (1) Processing the raw data, (2) Statistical analysis, and (3) Quality control (QC). The raw fastq files are submitted to esATAC and then undergo AdapterRemoval^28^ for adapter trimming and alignment to the human genome (GRCh38) using Bowtie2^29^. The mapped reads are sorted, duplicates are removed, and the reads are shifted for the Tn5 insertion^30^. esATAC identifies open chromatin peaks using F-seq31 which are then annotated, and gene ontology terms are summarized. QC reports are generated with raw sequencing read quality, fragment length QC analysis, and a number of QC plots to assess the quality of the reads and peak calling. Counts per peak were summarized using the featureCounts function from the subread package v.1.6.222. Read counts in the peak regions were then analyzed using the same procedure as for the Bulk RNAseq data. Only peaks in promoter regions were used for differential accessibility analyses.

### Gene set enrichment analysis

Functional gene set enrichment analysis (fgsea) was performed to explore the annotated genes or differential genes enriched hallmark pathways. For analysis of the transcriptomic heat shock conditions, fgsea was performed using hallmark pathways of the Molecular Signatures Database (MsigDB) and heat shock pathways in the Reactome Pathway Database (regulation of heat shock factor and HSF1-mediated heat shock response) and Gene Ontology Consortium (heat shock protein binding). For analysis of the transcriptomic thapsigargin conditions, fgsea was performed using a collection of pathways from the Gene Ontology Consortium, Reactome Pathway Database, and published gene sets^29,32^. For analysis of the epigenetic heat shock and thapsigargin conditions, fgsea was performed using transcription factor targets of the MsigDB. For all other conditions, the annotated genes or differential genes were analyzed based on the MsigDB to obtain all the involved hallmark pathway terms. The significant level of each pathway term was calculated by the Fisher test. Adjusted p-values < 0.05 were considered significant.

## Results

### Exposure to global ER stress via HS or thapsigargin results in stress- and cell type-specific transcriptional changes

As our group and others have identified ER stress to be a critical factor in the pathogenesis of ATTR and AL amyloidosis, we first sought to profile transcriptional signatures of cells exposed to HS and the SERCA kinase inhibitor thapsigargin. To accomplish this, immortalized neuronal SH-SY5Y and cardiac AC16 cells were subjected to 42°C for 110 minutes (HS) or thapsigargin (500 nM) for 16 hours **(Figure 1A)**. Exposure to HS resulted in marked gene expression changes in both SH-SY5Y and AC16 cells. SH-SY5Y cells exhibited differential expression of 7951 genes (3823 upregulated, 4128 downregulated), while AC16 cells exhibited 2818 differentially expressed genes (1350 upregulated, 1468 downregulated) **(Figure 1B, 1C)**. Top DEGs expressed by both SH-SY5Y and AC16 cells in response to HS included HSPA6, HSPA7, and HSPA1B as well as the proto-oncogene FOS **(Figure 1D)**.

**Figure 1.**
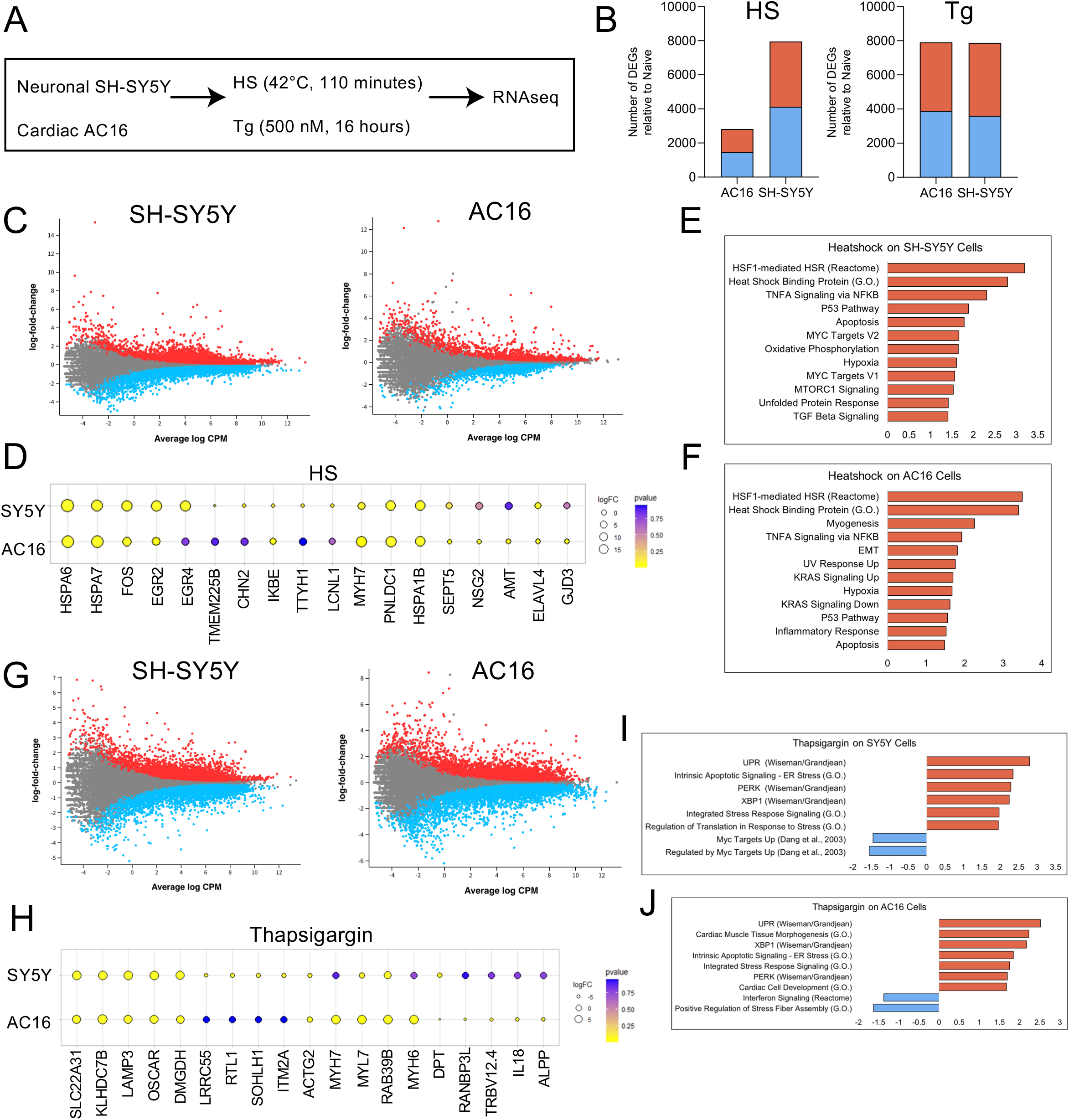
Cell type-specific differential gene expression of SH-SY5Y and AC16 cells in response to HS- and thapsigargin-induced ER stress. **(A)** Immortalized neuronal SH-SY5Y and cardiac AC16 cells were exposed to HS (42°C, 110 minutes) or thapsigargin (Tg) (500 nM, 16 hours). RNA was subsequently isolated from each sample and RNAseq was performed. Two technical replicates were used for each condition. **(B)** Number of differentially expressed genes (DEGs) relative to naïve (i.e. untreated cells) in each condition for SH-SY5Y and AC16 cells exposed to HS (left) or thapsigargin (right). Blue bars denote number of genes downregulated, red bars represent number of genes upregulated. FDR<0.05. **(C)** Differentially expressed genes in SH-SY5Y (left) and AC16 (right) cells exposed to HS. Each dot represents a single gene, red dots depict significantly upregulated genes, blue dots depict significantly downregulated genes. Grey dots depict genes which are not significantly changing. FDR<0.05. **(D)** Top upregulated genes in SH-SY5Y and AC16 cells upon exposure to heat shock. Yellow dots represent genes which are upregulated, blue dots represent genes which are downregulated. Size of dot denotes size of logFC. Color gradient represents adjusted p-value, yellow dots denote adjusted p-value < 0.05. **(E, F)** Gene ontology, reactome, and hallmark pathways represented in genes differentially expressed in SH-SY5Y **(E)** and AC16 **(F)** cells post-HS. **(G)** Differentially expressed genes in SH-SY5Y (left) and AC16 (right) cells exposed to thapsigargin. Red dots represent upregulated genes, blue dots represent downregulated genes, grey dots represent unchanging genes. FDR<0.05. **(H)** Top differentially expressed genes in SH-SY5Y and AC16 cells exposed to thapsigargin. Yellow represents upregulated genes, blue represents downregulated genes. Size of dot indicates magnitude of logFC. Color gradient represents adjusted p-value, yellow dots denote adjusted p-value < 0.05. **(I, J)** Published and curated gene sets represented in differentially expressed genes in SH-SY5Y **(I)** and AC16 **(J)** cells exposed to thapsigargin. Blue bars depict pathways represented in downregulated genes, red bars depict those represented in upregulated genes.

Unsurprisingly, pathway analysis revealed that SH-SY5Y and AC16 cells exposed to HS relate to known cellular responses to heat stress (e.g. the HSF1-mediated HSR Reactome pathway, Heat Shock Binding Protein GO pathway) and related TNFα Signaling (via NFKB hallmark pathway) **(Figure 1E, F)**. Interestingly, mTORC1 signaling pathway target genes were enriched in SH-SY5Y cells only, while KRAS signaling pathway genes were enriched in AC16 cells, suggesting potentially divergent downstream responses to HS for neuronal versus cardiac cells.

Exposure to toxic concentrations of thapsigargin resulted in differential expression of a similar number of genes in both SH-SY5Y (7880) and AC16 (7904) cells **(Figures 1B, 1G)**. SH-SY5Y and AC16 cells dosed with thapsigargin exhibited upregulation of several stress response pathways including: lysosomal pathway components (LAMP3), leukocyte receptor components (OSCAR), and mitochondrial matrix components (DMGDH) **(Figure 1H)**. As expected, UPR signaling pathway target genes (e.g. PERK, XBP1) were significantly enriched in upregulated genes for both SH-SY5Y and AC16 cells exposed to thapsigargin **(Figure 1J)**. Following 16 hour exposure exposure to 500nM thapsigargin, cells exhibited noticeable toxicity. In line with this noted toxicity, DEGs of both cell types were enriched for apoptotic signaling related to ER stress **(Figure 1I, J)**. Interestingly, SH-SY5Y cells downregulated gene sets associated with myc target genes while AC16 cells downregulated Interferon Signaling Reactome and Stress Fiber Assembly pathway target genes, again potentially suggesting divergent, cell type-specific response to stress **(Figure 1I, J)**.

### Cellular response to destabilized, disease-associated TTR and LC reveals cell type- and protein-specific transcriptional signatures

In an effort to profile the earliest instances of stress in response to disease-associated proteins such as TTR and LC, cells were exposed to bacterially-derived, recombinant human wild-type (TTR^WT^), neuropathy-associated (TTR^L55P^), and cardiomyopathy-associated (TTR^V122I^) TTR, as well as wild-type and destabilized AL LCs (AL^WT^ and AL^LC^, respectively) at physiological concentrations for 48 hours **(Figure 2A)**. Following dosing, RNA was isolated from each sample for bulk transcriptomic profiling via RNAseq. To control for differences due to the presence of misfolded TTRs, in parallel, recombinant TTRs were also pre-incubated with kinetic stabilizer tafamidis at an appropriate stoichiometric ratio. Although cells exposed to physiologically-relevant levels of wild-type and destabilized TTRs were morphologically unchanged, they exhibited many differentially expressed genes **(Figure 2B)**. Interestingly, neuronal SH-SY5Y cells exhibited a greater number of DEGs (relative to naïve cells) upon exposure to neuropathy-associated TTR^L55P^ than cardiomyopathy-associated TTR^V122I^ (3040 vs. 472, respectively) **(Figure 2B, left)**. Conversely, cardiac AC16 cells exhibited a slightly greater number of DEGs in response to cardiomyopathy-associated TTR^V122I^ than TTR^L55P^ (275 vs. 140, respectively) **(Figure 2B, right panel)**. Exposure to physiological concentrations of both pathologic and wild-type LCs resulted in marked increases in the number of DEGs when compared to recombinant TTR exposure. Lastly, both cell types exposed to wild-type TTR and LC exhibited a greater number DEGs compared to naïve (although less than their destabilized counterparts). As TTR and LC both have normal physiological processes (e.g. retinol transport and components of antibody structure, respectively), these DEGs could reflect how cells respond to said proteins under normal, homeostatic conditions. The top 10 up- and downregulated protein-coding genes in response to each stressor in SH-SY5Y and AC16 cells can be seen in **Tables 1** and **2**, respectively. For a comprehensive list of transcriptional changes please see **Data Files 1** and **2** for SH-SY5Y and AC16 cells, respectively.

**Figure 2.**
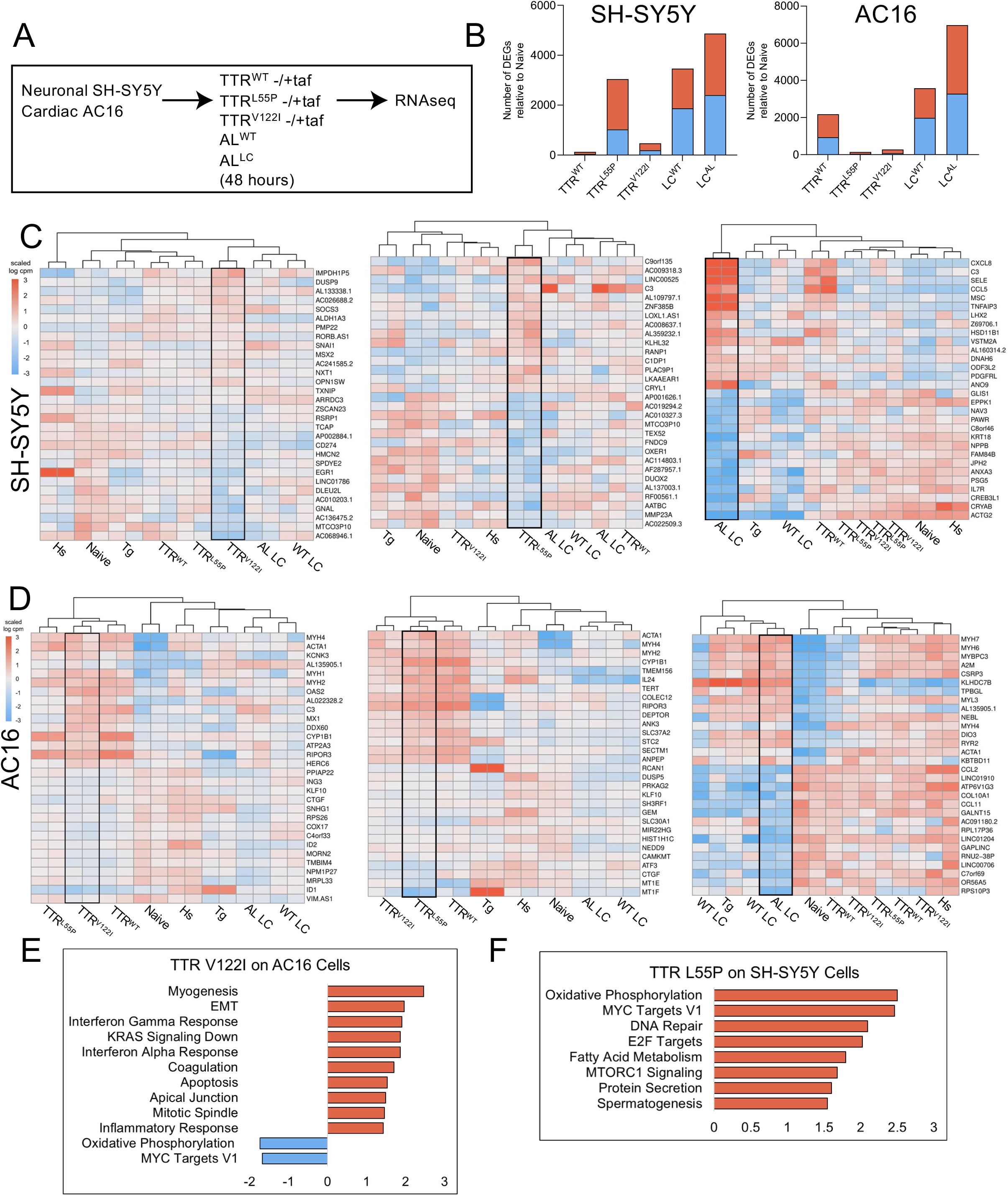
AC16 and SH-SY5Y cells exposed to amyloidogenic proteins exhibit robust cell type- and protein-specific gene expression changes. **(A)** SH-SY5Y and AC16 cells were exposed to bacterial-derived recombinant human TTR (wild-type, neuropathy-associated TTR^L55P^, cardiomyopathy-associated TTR^V122I^) and (wild-type, AL^WT^, and destabilized, AL^LC^) variants at physiologically relevant concentrations for 48 hours. To control for gene expression changes related to misfolded forms of TTRs, in parallel, TTRs were pre-incubated with kinetic stabilized tafamidis at appropriate stoichiometric ratios. RNA was isolated from each condition for RNAseq. Two technical replicates were used for each condition. **(B)** Number of differentially expressed genes (DEGs) relative to naïve (i.e. untreated cells) in each condition for SH-SY5Y (left) and AC16 (right) cells. Blue bars denote number of genes downregulated, red bars represent number of genes upregulated. FDR<0.05. **(C, D)** Top 15 significantly up- and downregulated genes seen in SH-SY5Y **(C)** and AC16 **(D)** cells exposed to TTRs, LCs, HS, and thapsigargin. Each individual heatmap depicts the top 30 differentially expressed genes for cardiomyopathy-associated TTR^V122I^ (left), neuropathy-associated TTR^L55P^ (middle), or destabilized LC. Fold-changes are determined relative to naïve (i.e. untreated cells). Stimuli are depicted on the x-axis, differentially expressed genes on the y-axis. Samples are arranged via hierarchical clustering. FDR<0.05. **(E, F)** Hallmark pathways enriched in SH-SY5Y cells exposed to neuropathy-associated TTR^L55P^ **(E)** and AC16 cells exposed to cardiomyopathy-associated TTR^V122I^ **(F)**. Blue bars depict pathways enriched in downregulated genes, red pathways depict those enriched in downregulated genes.

**Table 1.**
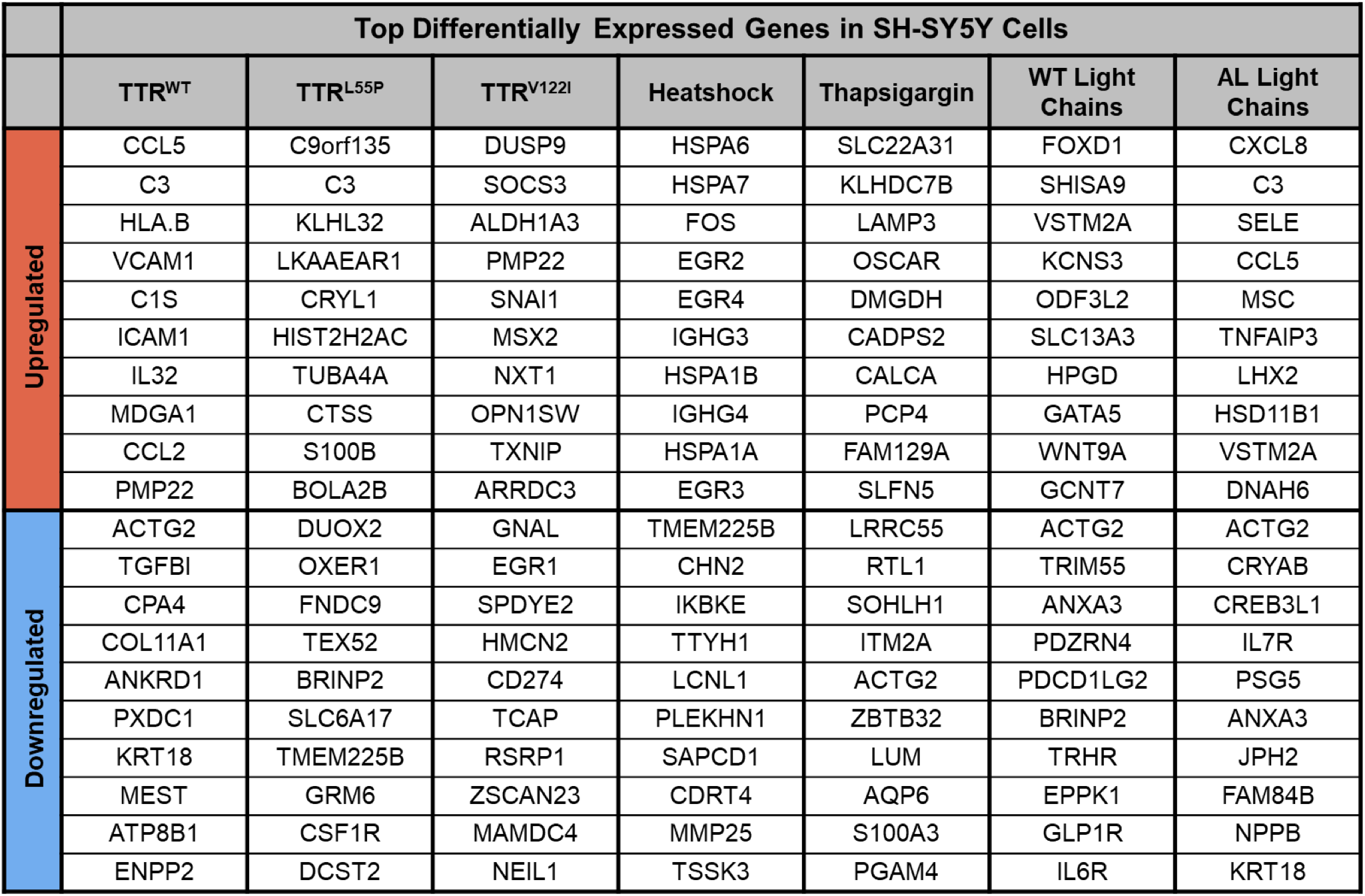
Top 10 up- and top 10 downregulated genes in neuronal SH-SY5Y cells exposed to HS, thapsigargin, or destabilized proteins.

**Table 2.** Top 10 up- and top 10 downregulated genes in cardiac AC16 cells exposed to HS, thapsigargin, or destabilized proteins.

| Top Differentially Expressed Genes in AC16 Cells |  |  |  |  |  |  |  |
| --- | --- | --- | --- | --- | --- | --- | --- |
|  | TTR<sup>WT</sup> | TTR<sup>L55P</sup> | TTR<sup>V122I</sup> | Heatshock | Thapsigargin | WT Light Chains | AL Light Chains |
| Upregulated | MYH4 | ACTA1 | MYH4 | HSPA6 | KLHDC7B | KLHDC7B | MYH7 |
|  | ACTA1 | MYH4 | ACTA1 | HSPA7 | MYH7 | TPBGL | MYL7 |
|  | KCNK3 | MYH2 | KCNK3 | MYH7 | MYL7 | ESRP1 | MYH6 |
|  | STK32A | CYP1B1 | MYH1 | PNLDC1 | RAB39B | ACTA1 | PLN |
|  | CALML4 | TMEM156 | MYH2 | HSPA1B | MYH6 | BRINP3 | MYL2 |
|  | VIPR1 | SERPINB2 | OAS2 | MYH6 | PLN | FEZ1 | TNNI1 |
|  | PKNOX2 | TMEM119 | C3 | HSPA1A | THBD | GALR2 | MYBPC3 |
|  | MYH1 | IL24 | MX1 | MYBPC3 | TNNI1 | DEGS2 | A2M |
|  | MYH2 | TERT | DDX60 | RBFOX3 | SLC22A31 | BEX2 | CSRP3 |
|  | CYP1B1 | COLEC12 | ISG20 | NHLH1 | MT1G | ST6GALNAC3 | KLHDC7B |
| Downregulated | STMN2 | ELAVL4 | STMN2 | SEPT5 | DPT | ATP6V1G3 | GPR183 |
|  | CDH26 | PRAME | ELAVL4 | NSG2 | RANBP3L | HAPLN1 | OR56A5 |
|  | NSG2 | MT1F | PRAME | AMT | TRBV12.4 | TRBV12.4 | JCHAIN |
|  | PLXNA4 | CNTN1 | CNTN1 | ELAVL4 | IL18 | CCL11 | TNFSF18 |
|  | ELAVL4 | PTPRQ | RPS29 | GJD3 | ALPP | PLP1 | PTPRC |
|  | EML5 | TNFSF4 | RPL22L1 | HOXD4 | CLEC3B | COL10A1 | IL26 |
|  | ANKRD6 | MT1E | SNRPG | ARFGEF3 | AKR1C3 | SCN3A | GALNT15 |
|  | PRAME | CTGF | ID1 | CCDC39 | TRBV12.3 | GALNT15 | LAD1 |
|  | SCT | WNT2B | RPS21 | PRAME | SCN3A | PSG1 | CCL11 |
|  | SLC6A2 | RGS5 | MRPL33 | CLUL1 | PLP1 | PSG2 | LYPD6B |

Comparisons of the top 30 DEGs (top 15 upregulated and top 15 downregulated) upon dosing SH-SY5Y cells with neuropathy-associated TTR^L55P^, cardiomyopathy-associated TTR^V122I^, and mutant LC highlighted transcriptional signatures distinct to each respective stressor **(Figure 2C)**. Interestingly, the transcriptional signature elicited by neuropathy-associated TTR^L55P^ may be more robust than that elicited by cardiomyopathy-associated TTR^V122I^. Moreover, AC16 cells exposed to TTR^V122I^ potentially exhibit a unique transcriptional signature that is more robust than AC16 cells exposed to TTR^L55P^ albeit to a slightly lesser degree **(Figure 2D)**. As reflected in the number of DEGs in response to AL^LC^, destabilized LC exposure resulted in strong transcriptional signatures in both SH-SY5Y and AC16 cells **(Figure 2C, 2D)**.

Gene set enrichment analysis revealed upregulation of diverse hallmark pathways in both mutant TTR and LC conditions **(Figure 2E, 2F, S1)**. Neuronal cells exposed to both TTR^L55P^ and TTR^V122I^ exhibited upregulation of genes associated with DNA repair, E2F targets, mTORC1 signaling, Myc targets, and oxidative phosphorylation **(Figure 2E, S1)**, potentially highlighting a generalized SH-SY5Y stress response program. Upregulation of fatty acid metabolism and protein secretion associated genes were specific to TTR^V122I^ exposure, while upregulation of the complement system was specific to TTR^L55P^ exposure in SH-SY5Y cells. Gene set enrichment analysis further revealed enrichment of diverse hallmark pathways in cardiac cells dosed with amyloidogenic protein **(Figure 2F, S1)**. Upon both TTR^L55P^ and TTR^V122I^ exposure, DEGs in cardiac cells were associated with upregulation for myogenesis and coagulation and downregulation for Myc targets. AC16 cells exposed to TTR^L55P^ showed upregulation for epithelial-mesenchymal transition, interferon gamma and alpha response, KRAS activation, apoptosis, apical junction complex, mitotic spindle assembly, and inflammatory response. AL^LC^ exposure in SH-SY5Y cells showed upregulation of genes associated with many pathways including: Myc targets, oxidative phosphorylation, DNA repair, reactive oxygen species pathway, inflammatory response, interferon gamma response, and mTORC1 signaling.

### SH-SY5Y and AC16 cells exposed to HS, thapsigargin, and destabilized proteins exhibit differential expression of a subset of overlapping genes

In addition to comparing how neuronal and cardiac cells respond to pathologically-diverse TTRs and LCs, we next sought to compare the signature of these stressors with that following exposure to HS- and thapsigargin-induced stress. To accomplish this, we compared DEGs of SH-SY5Y and AC16 cells exposed to HS and thapsigargin to those exposed to destabilized TTR and LC.

SH-SY5Y cells subjected to mutant TTRs, HS, and thapsigargin had 135 DEGs in common **(Figure 3A)**, while those exposed to destabilized LC, HS, and thapsigargin differentially expressed 720 shared genes **(Figure 3B)**. SH-SY5Y cells exposed to destabilized TTRs and LCs shared differential expression of 85 genes **(Figure 3C)**. Perhaps due in part to lower numbers of DEGs upon stimulus exposure, in contrast, AC16 cells dosed with TTRs or HS shared differential expression of 22 genes **(Figure 3D)**. Similarly, AC16 cells exposed to HS, thapsigargin, or ALLC differentially expressed 339 common genes **(Figure 3E)**. Lastly, AC16 cells exposed to amyloidogenic TTRs or AL^LC^ differentially expressed only 10 common genes **(Figure 3F)**, suggesting activation of unique transcriptional stress response pathways.

**Figure 3.**
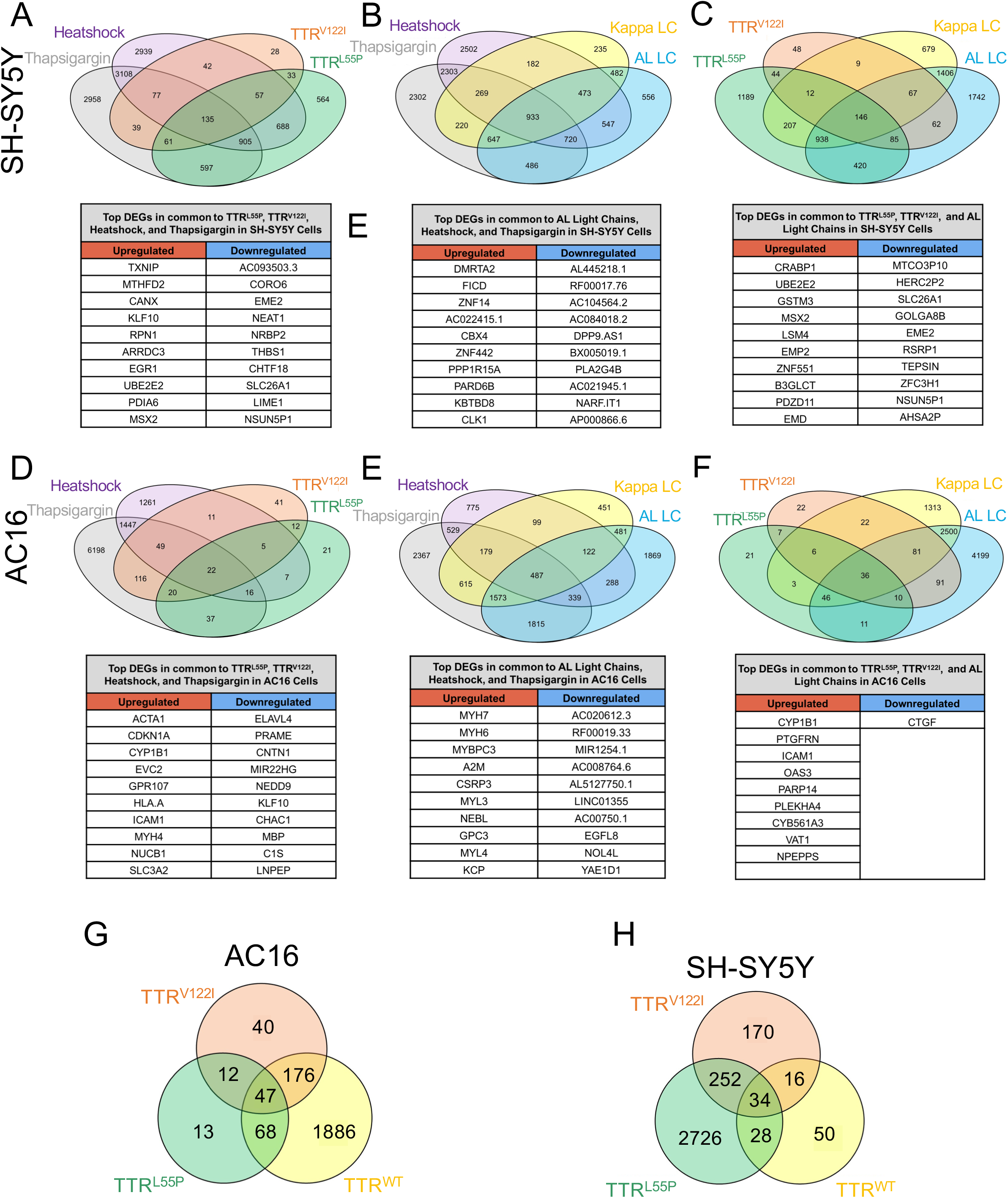
Cells exposed to destabilized TTR and LC exhibit gene expression changes distinct and shared with those subjected to HS- and thapsigargin-induced ER stress. **(A-C)** Differentially expressed genes shared between SH-SY5Y cells exposed to HS, thapsigargin, cardiomyopathy-associated TTR^V122I^, neuropathy-associated TTR^L55P^, as well as wild-type and destabilized LC (AL^LC^). For each comparison, the top 10 DEGs up- and downregulated in common to all comparisons are listed below. **(D-F)** DEGs shared across AC16 cells exposed to HS, thapsigargin, TTRV122I, TTRL55P, and wild-type and AL LCs. Top 10 DEGs in common to all conditions are listed below. **(G, H)** Differentially expressed genes broken down by TTR variant in exposed AC16 **(G)** and SH-SY5Y **(H)** cells. Top 5 upregulated genes specific to TTR^V122I^ **(G)** and TTR^L55P^ **(H)** are noted.

As reflected in **Figure 3**, in both SH-SY5Y and AC16 cells, neuropathy- and cardiomyopathy-associated TTRs drove differential expression of genes unique to each variant. In AC16 cells, for example, TTR^V122I^ elicited differential expression of 40 unique genes **(Figure 3G)**, while in SH-SY5Y cells, TTR^L55P^ exposure resulted in differential expression of 2726 unique genes **(Figure 3H)**.

### Neuronal and cardiac cells exhibit changes in chromatin accessibility profiles in response to HS and thapsigargin exposure

We next wanted to expand upon our transcriptomic profiles of diverse cellular stress by assessing the effect of each stimuli at the chromatin level. To this end, we employed ATACseq to investigate changes in chromatin organization following exposure to HS and thapsigargin **(outlined in Figure 4A)**. In doing so, we found that HS and thapsigargin resulted in robust changes in differentially accessible peaks (DAPs) located within promoter regions of genes in both SH-SY5Y and AC16 cells **(Figure 4B)**.

**Figure 4.**
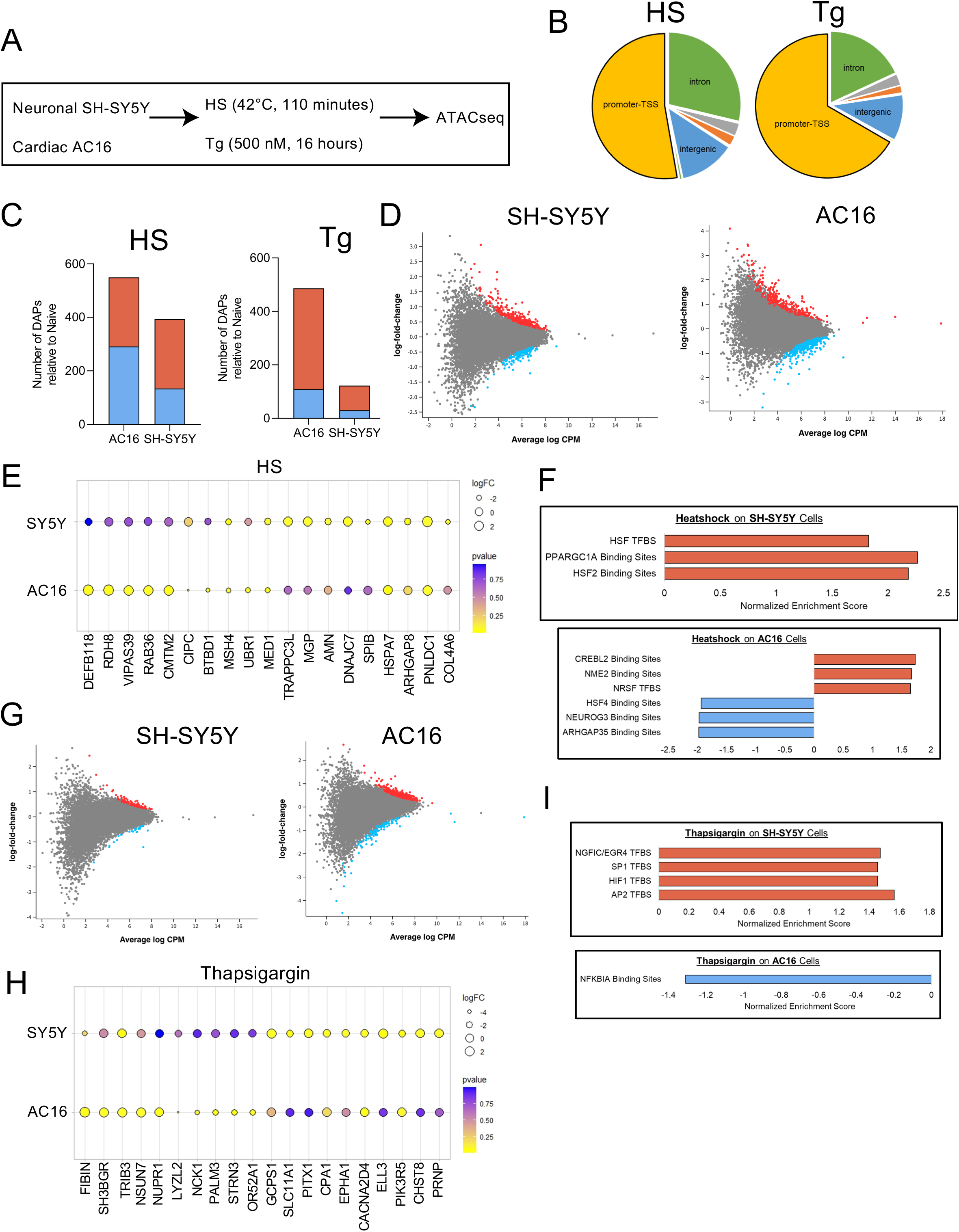
SH-SY5Y and AC16 cells exposed to HS and thapsigargin exhibit changes in chromatin accessibility signatures in cell type- and stress-specific manners. **(A)** SH-SY5Y and AC16 cells were exposed to HS (42°C, 110 minutes) or thapsigargin (500 nM, 16 hours). **(B)** Percentage of total DAPs in AC16 cells exposed to HS or thapsigargin located within promoter regions, intergenic regions, intronic regions, or other non-coding regions of the genome. **(C)** Number of differentially accessible peaks (DAPs) in SH-SY5Y and AC16 cells exposed to HS and thapsigargin (Tg), left and right respectively. DAPs are relative to naïve (i.e. untreated cells). DAPs included in calculation represent those associated with the promoter regions of known protein coding genes as defined by distance to the transcriptional start site. FDR<0.05. Red represents DAPs which increase in abundance post-stress, while blue represents DAPs which decrease post-stress. **(D)** Significant DAPs in SH-SY5Y (left) and AC16 (right) cells exposed to heat shock. Red dots depict DAPs which increase in abundance, blue denotes DAPs which decrease in abundance. Fold-change relative to naïve condition. FDR<0.05. **(E)** Top DAPs associated with promoter regions whose accessibility changes in SH-SY5Y and AC16 cells exposed to HS. Size of dot represents strength of fold-change. Color gradient represents adjusted p-value. **(F)** Transcription factor binding sites associated with DAPs in promoter regions post-HS in SH-SY5Y (top) and AC16 (bottom) cells. Red bars depict transcription factor targets enriched in DAPs becoming more accessible post-HS while blue bars represent pathways enriched in DAPs becoming less accessible post-HS. **(G)** Total number of promoter region-associated DAPs differentially accessible in SH-SY5Y (left) and AC16 (right) cells post-thapsigargin exposure. Red and blue dots depict increasing and decreasing DAPs relative to naïve cells, respectively. FDR<0.05. **(H)** Top changing promoter-associated DAPs in SH-SY5Y and AC16 cells exposed to thapsigargin. **(I)** Transcription factor binding sites associated with DAPs in promoter regions post-thapsigargin exposure in SH-SY5Y (top) and AC16 (bottom) cells.

A majority of newly formed DAPs resulting in response to both HS and thapsigargin as compared to naïve cells resided in promoter regions (as defined by distance to the transcriptional start site), suggesting changes in chromatin accessibility could result in changes in expression of protein-coding genes **(Figure 4C, S2)**. Based on these data, unless otherwise stated, all ATACseq data listed below focuses on DAPs found in promoter regions of protein-coding and noncoding genes. The top 10 up- and downregulated DAPs found within promoter regions of SH-SY5Y and AC16 cells in response to each stressor can be found in **Table 3**. For a comprehensive list of DAPs occurring in stressed SH-SY5Y and AC16 cells see **Data Files 3** and **4**, respectively.

**Table 3.** Top 10 up- and top 10 downregulated DAPs in AC16 and SH-SY5Y cells exposed to HS and thapsigargin.

|  | Top Differentially Accessible Peaks |  |  |  |
| --- | --- | --- | --- | --- |
|  | AC16 Cells |  | SH-SY5Y Cells |  |
|  | Heatshock | Thapsigargin | Heatshock | Thapsigargin |
| Upregulated | MIR23B | FIBIN | PNLDC1 | SNORA80D |
|  | DEFB118 | SH3BGR | KCNMB2-AS1 | ELL3 |
|  | LINC00310 | LINC00824 | MIR548T | GGPS1 |
|  | NAPSB | MYHAS | MASCRNA | C1orf158 |
|  | RDH8 | TRIB3 | MIR4647 | PRNP |
|  | MIR3163 | NSUN7 | HSPA7 | PITX1 |
|  | LOC101928401 | LINC0056 | LOC100129697 | LINC02593 |
|  | MIR6501 | LINC01686 | LINC00251 | CHST8 |
|  | LINC00996 | LOC105371267 | LOC101928540 | MAN1B1-DT |
|  | VIPAS39 | NUPR1 | TRAPPC3L | RAD23A |
| Downregulated | CIPC | LYZL2 | SPIB | SLC11A1 |
|  | BTBD1 | ZNF775 | COL4A6 | ROCK1P1 |
|  | MSH4 | NCK1 | MSH4 | PIK3R5 |
|  | UBR1 | MIR6763 | MIR5193 | CPA1 |
|  | RNU5A-1 | PALM3 | ARHGAP8 | CACNA2D4 |
|  | MED1 | STRN3 | ZNF140 | EPHA1 |
|  | CCDC22 | OR52A1 | EWSAT1 | NUP43 |
|  | SYTL4 | MYOT | AMN | MRPL13 |
|  | RNU11 | FXYD3 | RNU5A-1 | SART3 |
|  | ZNF226 | LINC00303 | CACNA2D4 | LINC02623 |

**Table 4.** Top 10 up- and top 10 downregulated DAPs in AC16 cells exposed to TTRWT and mutant TTRs.

|  | Top Differentially Accessible Peaks in AC16 Cells |  |  |
| --- | --- | --- | --- |
|  | TTR<sup>WT</sup> | TTR<sup>L55P</sup> | TTR<sup>V122I</sup> |
| Upregulated | LRRN4CL | SNORA74C-1 | SND1-IT1 |
|  | C9orf170 | LOC102724050 | IDH2 |
|  | C1QTNF1-AS1 | C1QTNF1-AS1 | LRRN4CL |
|  | FRRS1 | EGOT | LINC02323 |
|  | LOC100132057 | LRRN4CL | C1QTNF1-AS1 |
|  | BEND6 | RPL14 | RNVU1-6 |
|  | SLC16A4 | C9orf170 | RPL14 |
|  | IDH2 | BEND6 | LINC02175 |
|  | NUPR1 | RPL13AP20 | C9orf170 |
|  | RPL14 | RNVU1-6 | LINC00111 |
| Downregulated | MIR4476 | SNRPN | LINC02313 |
|  | LINC02067 | GPATCH3 | ZCWPW2 |
|  | LINC01826 | LOC105378305 | MIR328 |
|  | MIR148A | STIMATE | MIR3176 |
|  | LINC01170 | LINC00613 | PINLYP |
|  | ALG5 | CACNA1C-IT2 | LELP1 |
|  | SCGB1D1 | FAM71F2 | PDE6C |
|  | ERP27 | LINC02624 | PHKA2-AS1 |
|  | MIR4655 | ELFN1 | TSSK2 |
|  | PRR27 | LINC01826 | LOC729558 |

In both SH-SY5Y and AC16 cells, HS resulted in many changes in chromatin accessibility in the form of DAPs (393 for SH-SY5Y cells and 549 for AC16 cells **(Figures 4B, 4D)**. Some of the DAPs with greatest effects post- HS in both SH-SY5Y and AC16 cells include those associated with genes involved in HS (HSPA7), mRNA processing (PNLDC1), and rhoGTPase activity (ARHGAP8), potentially suggesting functional outcomes resulting from HS-induced chromatin accessibility changes **(Figure 4E, 4F)**. In SH-SY5Y cells, HS exposure resulted in DAPs forming in HSF transcription factor binding sites as well as PPARGC1A (master regulator of mitochondrial biogenesis) **(Figure 4F)**. In AC16 cells, HS exposure resulted in increased accessibility of CREB-like binding sequences (CREBL2) and decreased accessibility of HSF4 binding sequences **(Figure 4F)**.

Despite observed toxicity, thapsigargin resulted in fewer DAPs in both SH-SY5Y and AC16 cells as compared to HS (122 for SH-SY5Y and 486 for AC16 cells) **(Figures 4B, 4G)**. For both SH-SY5Y and AC16 cells, top DAPs increasing in accessibility upon thapsigargin exposure included those lying in the promoter regions of genes associated with PI3K signaling (PIK3R5) and pro-apoptotic pathways (TRIB3) **(Figure 4H)**. SH-SY5Y cells exposed to thapsigargin exhibited increased accessibility in DAPs associated with proliferation-promoting transcription factor binding sites (EGR4), regulation of apoptosis and post-transcriptional modifications (SP1), and response to hypoxia (HIF1) **(Figure 4I)**. Alternatively, AC16 cells exhibited decreased accessibility in immune-regulating NFKBIA binding sites **(Figure 4I)**.

### Destabilized TTR exposure results in variant- and cell type-specific chromatin accessibility signatures that are potentially resolved by pre-incubation with kinetic stabilizer

As our above data suggest that neuronal and cardiac cells experience chromatin level changes in response to HS- and thapsigargin-mediated stress, we next wanted to extend this experiment and analysis to investigate chromatin level changes in response to destabilized, disease-causing TTRs. To accomplish this, SH-SY5Y and AC16 cells were again exposed to physiologically-relevant concentrations of recombinant human TTR^WT^, neuropathy-associated TTR^L55P^, or cardiomyopathy-associated TTR^V122I^. After exposure to TTRs for 48 hours, gDNA was isolated from each condition and ATACseq was performed. In parallel, in an effort to control for changes resulting from exposure to misfolded forms of each protein, each TTR was pre-incubated with the kinetic stabilizer tafamidis at an appropriate stoichiometric ratio.

By PCA, as expected, the greatest differences in chromatin accessibility were dictated by cell type **(Figure S2A, left)**. Additionally, as with our transcriptomic data, for both SH-SY5Y and AC16 cells, HS and thapsigargin exhibited the greatest changes when compared to naïve cells, forming distinct groups by PCA **(Figure S2A, middle, right)**. As with HS- and thapsigargin-exposed cell most DAPs observed in AC16 cells dosed with TTRs lie in regions associated with promoters and the transcriptional start site, followed by intronic and intergenic regions **(Figure S2B)**.

Although we sought to compare chromatin level changes in response to disease-causing, misfolding-prone TTRs to HS and thapsigargin, in doing so, we found that DAP signatures appear to be less similar across stressors than transcriptional signatures, as evidenced by only 8 DAPs being common across HS, thapsigargin, TTR^V122I^, and TTR^L55P^ **(Figure S2C)**. Despite this discordance, we then wanted to determine differences in DAPs resulting from exposure to different TTR species. As AC16 cells exhibited a greater response to TTRs at the DAP level as compared to SH-SY5Y, in these analyses we focused on TTR-exposed AC16 cells. To this end, AC16 cells share 93 DAPs in common when exposed to TTR^L55P^ or TTR^V122I^, demonstrating a partially shared chromatin-level response to destabilized TTRs **(Figure 5B)**. As with the aforementioned transcriptional data, AC16 cells exposed to TTR^L55P^ and TTR^V122I^ appear to exhibit variant-specific responses at the level of chromatin accessibility.

**Figure 5.**
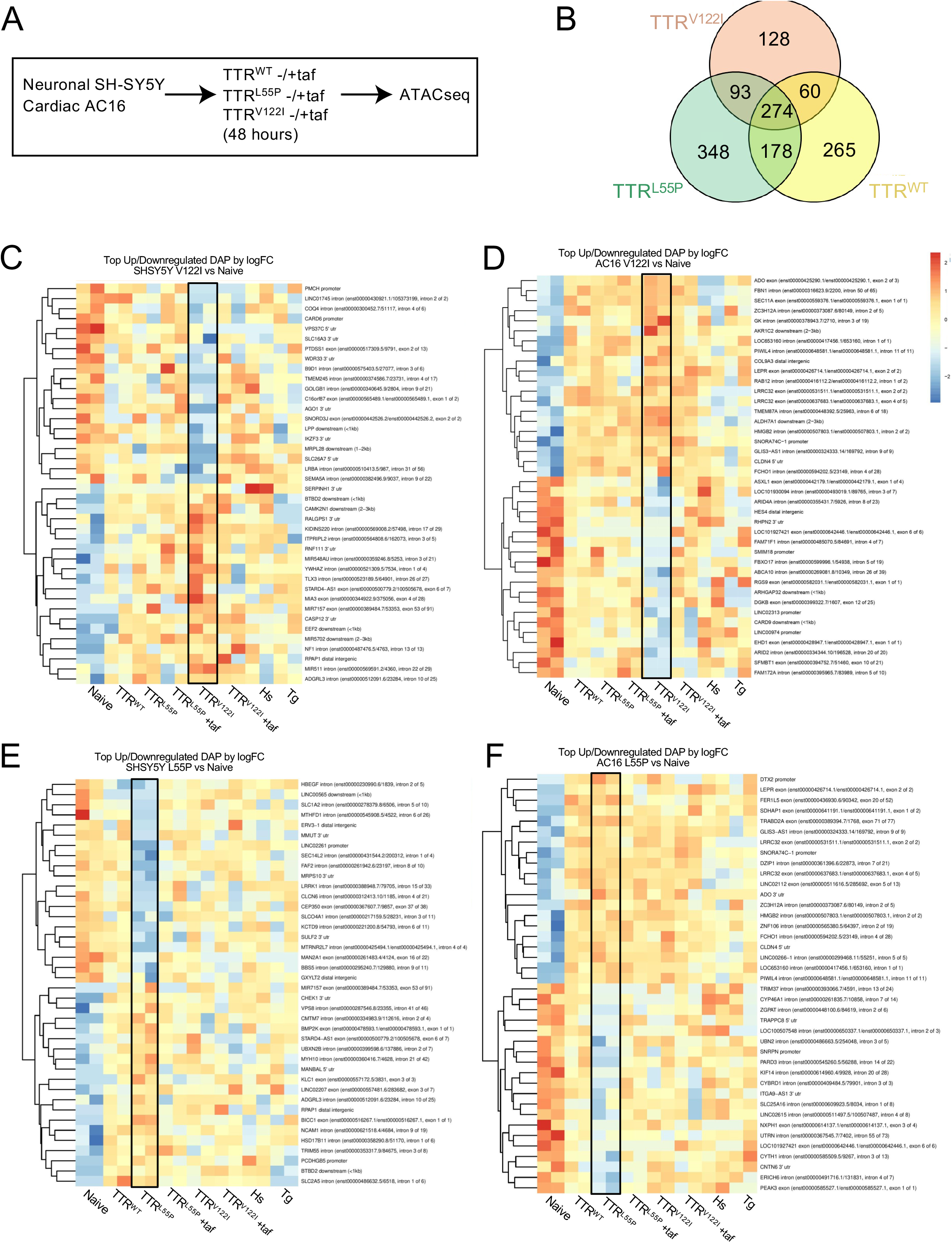
Cells exposed to pathologically-associated destabilized TTR exhibit mutation-specific changes in chromatin signatures resolved by pre-incubation with tafamidis. **(A)** SH-SY5Y and AC16 cells were exposed to TTR^WT^, neuropathy-associated TTR^L55P^, and cardiomyopathy-associated TTRV122I at physiological concentrations (0.2 mg/mL). 48 hours later, gDNA was isolated and ATACseq was performed. In parallel, recombinant proteins were pre-incubated with tafamidis (taf) to control for changes due to misfolded forms of the protein. **(B)** Comparison of promoter region-associated DAPs unique to and shared between AC16 cells exposed to wild-type (TTR^WT^), cardiomyopathy- (TTR^V122I^), or neuropathy-associated (TTR^L55P^) TTR variant. DAPs noted are those relative to naïve condition. **(C-F)** Heatmaps depicting top 20 up- and downregulated DAPs for SH-SY5Y and AC16 cells exposed to cardiomyopathy-associated TTR^V122I^ **(C, D)** and neuropathy-associated TTR^L55P^ **(E, F)** in the absence of tafamidis. On the y-axis are DAPs occurring in promoter regions associated with protein coding genes for each noted variant relative to naïve condition. HS- and Tg-exposed samples (noted in **Fig. 4**) are also included. The x-axis notes samples in technical duplicate. Black boxes depict DAP signature resulting from noted variant.

Lastly, we then wanted to visualize the strongest DAP signature driven by each pathologic variant across both SH-SY5Y and AC16 cells. To do this, we examined the top 20 up and downregulated DAPs in SH-SY5Y and AC16 cells exposed to TTRs in the presence and absence of tafamidis. When portraying these data via heatmaps, we see clear signatures denoting both increased and decreased accessibility of DAPs lying within promoter regions of protein coding genes **(black boxes, Figure 5C-F)**. Upon pre-incubating TTR^L55P^ and TTR^V122I^ with tafamidis immediately prior to dosing cells, these signatures appear to resolve, and accessibility of top DAPs return to levels comparable to cells dosed with TTR^WT^, suggesting that said changes are due to the misfolded form of the dosed TTR.

## Discussion

Here, we mapped transcriptional and chromatin level changes resulting from exposure to profound ER stress via HS and thapsigargin or disease-associated, amyloidogenic TTR and LC proteins.

As our group and others have identified differential regulation of ER stress machinery in cells producing and interacting with pathology-associated TTRs and LCs^10,11^, we sought to compare these responses to changes elicited by HS and thapsigargin. In doing so, we generated transcriptional and chromatin accessibility profiles of neuronal SH-SY5Y and cardiac AC16 cells undergoing HS and global UPR activation via thapsigargin. Although transcriptional responses to these perturbagens have been well-documented, these resources are largely generated in single cell types; our data however, overlays cell type-specific responses to these widely used conditions. Additionally, through our ATACseq experiment, we provide evidence that exposure to ER stress (via HS and thapsigargin) results in substantial changes in accessibility of chromatin regions throughout the genome that are specific to each cell type. To this end, interestingly, post-HS, SH-SY5Y cells exhibited increased accessibility of chromatin regions located within transcription factor binding sites for HSF1, perhaps reflecting functional consequences of said changes in contributing to HSR.

The systemic amyloid diseases are a class of highly complex disorders involving the production of proteins which misfold, travel throughout circulation, and deposit as proteotoxic aggregates at distal target organs. Despite advances in a number of therapeutics for patients with these diseases, the mechanisms of how downstream target cell types become damaged by protein aggregates remain unclear. Here, we employed RNAseq and ATACseq to profile molecular responses of neuronal and cardiac cells to bacterially-derived, destabilized recombinant human TTR and LC. Importantly, as patients with ATTR and AL amyloidosis are typically diagnosed in late adulthood (coinciding with prolonged accumulation of misfolded protein)^4–6^, our experiments map the earliest signatures of cellular stress resulting from destabilized protein exposure and could potentially lead to the identification of novel prognostic biomarkers.

At the transcriptional level, neuronal SH-SY5Y and cardiac AC16 cells appeared to respond more strongly to their corresponding pathologically-associated TTR, as implied by a greater number of DEGs. Interestingly, this observation was not seen at the chromatin level, suggesting that mutation-driven organ tropism could be reflected at the level of transcriptional stress response. As SH-SY5Y and AC16 cells both responded to each TTR in unique ways, attention should be paid to differences between each TTR variant. To this end, recent work has identified several structural differences across different TTR mutants^33–35^. Although all TTRs employed in this study were produced via bacteria and thus lack post-translational modifications, said proteins could adopt various conformations when in contact with the extracellular environment of each cell type and thus elicit novel responses.

Through our RNAseq data, we observed that AC16 cells exposed to TTR^V122I^, TTR^L55P^, and AL^LC^ exhibited upregulation of myogenesis Hallmark pathway target genes, potentially representing a protective mechanism (e.g. cardiac remodeling in instances of stress). Interestingly, patients with both ATTR and AL amyloidosis exhibit symptoms associated with cardiac hypertrophy^36,37^ which is reflected in our transcriptional signatures. At the same time, SH-SY5Y cells exposed to TTR^V122I^, TTR^L55P^, and AL^LC^ exhibited increased expression of genes associated with mTORC1 signaling, E2F targets, and oxidative phosphorylation Hallmark pathways **(Figure 3C, D)**. Future efforts may seek to utilize pharmacologic effectors of mTOR signaling to modify destabilized protein-mediated stress. While pathways similarly upregulated across variants could reflect generalized stress responses, discordant signatures may represent novel responses specific to each amyloidogenic protein or mutant. For example, SH-SY5Y cells exposed to TTR^L55P^ resulted in increased expression of genes associated with fatty acid metabolism and protein secretion, both processes implicated in the pathogenesis of other amyloid disorders such as Alzheimer’s Disease (AD)^35,36^. Taking this into consideration, clinical data from patients with ATTR-associated polyneuropathy may uncover the potential of circulating metabolites as novel biomarkers for identifying onset of ATTR amyloidosis as well as other amyloid disorders with similar mechanisms of action (e.g. AD, etc.).

In addition to transcriptional responses, our data demonstrate that cells interacting with extracellular TTRs exhibit chromatin-level changes. As DAP signatures appear to resolve upon pre-incubating destabilized variants with tafamidis, said changes may result from misfolded forms of the administered protein. Although we did not observe changes at the transcriptional level when pre-incubating TTRs with tafamidis (perhaps due to dose, exposure, and/or timing), these results highlight the benefit of structure-based kinetic stabilizers at downstream target cell types.

A majority of current treatments for ATTR amyloidosis as well as pre-clinical pharmacologic agents for AL amyloidosis primarily focus on decreasing levels of circulating misfolded protein, thus limiting the formation of downstream toxic aggregates ^40–42^. Understanding the molecular mechanisms through which TTRs and LCs elicit damage however, may uncover novel therapeutic avenues applicable at downstream target cells. At the same time, efficacy of treatments for systemic amyloid diseases depends on disease progression – a notion complicated by a lack of biomarkers for these difficult to diagnose diseases. Future work may seek to harness these data to identify diagnostic biomarkers indicating the earliest instances of TTR- and LC-mediated stress. Through these efforts, we have generated comprehensive datasets representing transcriptional- and chromatin level responses of SH-SY5Y and AC16 cells to ER stress via HS and thapsigargin as well as amyloidogenic TTRs and LCs. In doing so, we identified transcriptional signatures of TTR- and LC-mediated cell stress unique to both cell type and destabilized protein. In addition, our analysis uncovers changes in chromatin architecture resulting from the presence of mutant TTR deposition. These changes were absent following pre-incubation with tafamidis, suggesting that the stabilizer may prevent potentially detrimental changes observed at the chromatin level. The datasets generated herein may be utilized by the proteostasis and amyloid research communities to better understand ER stress in multiple cellular contexts as well as aid in the development of cellular models for complex, multi-system amyloid diseases such as ATTR and AL amyloidosis.

## Acknowledgments

The authors thank the Wiseman laboratory of The Scripps Research Institute for providing recombinant human TTRs and tafamidis, the Microarray and Sequencing Resource Core Facility of Boston University School of Medicine for technical assistance with sample preparation, Dr. Gregory J. Miller of the Center for Regenerative Medicine (CReM) for technical and operational support, and Dr. Kim Vanuytsel and Todd Dowrey of the CReM for helpful discussions. This work was supported by the National Institutes of Health - NIDDK (grants R01DK102635 [G.J.M.], F31DK121481 [R.M.G.]), NCATS (grant 1UL1TR001430 [R.M.G., G.J.M.]), the Amyloidosis Foundation, and the Young Family Amyloid Research Fund.

## Author Contributions

S.G., R.M.G., and G.J.M. designed the project, devised experiments, analyzed data, and wrote the manuscript. R.M.G., S.G., C.V.M., J.L.V., D.K., C.S.G., and C.V.E. performed experiments and analyzed data. V. S. and L.H.C. provided feedback and assisted in writing the manuscript.

---

**Supplemental Figure 1.**
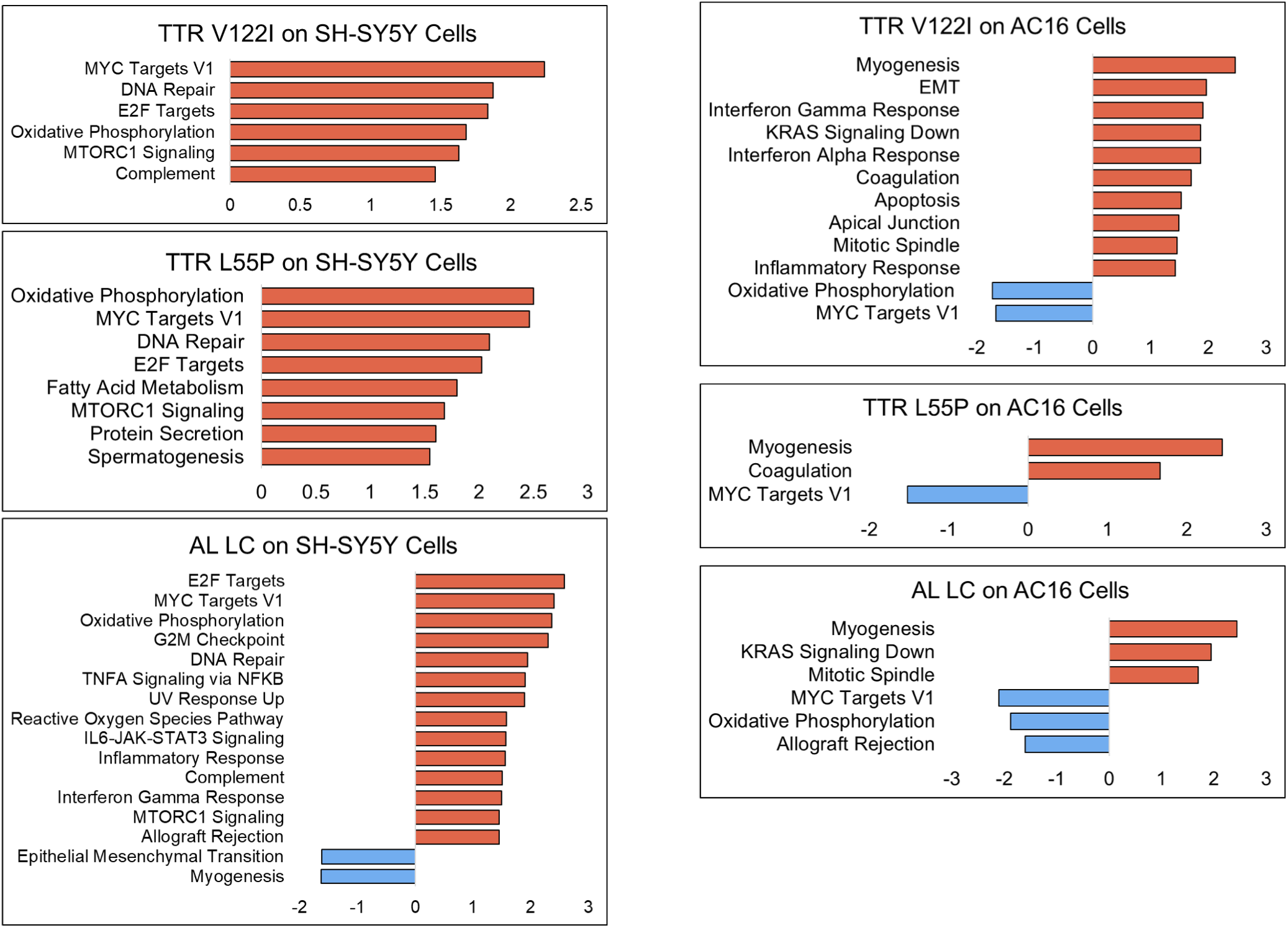
Pathway analysis of SH-SY5Y and AC16 cells exposed to pathologically-diverse amyloidogenic TTR and AL LC. DEGs (FDR<0.05) resulting from exposure of SH-SY5Y and AC16 cells to destabilized TTRs and LCs were subjected to pathway analysis.

**Supplemental Figure 2.**
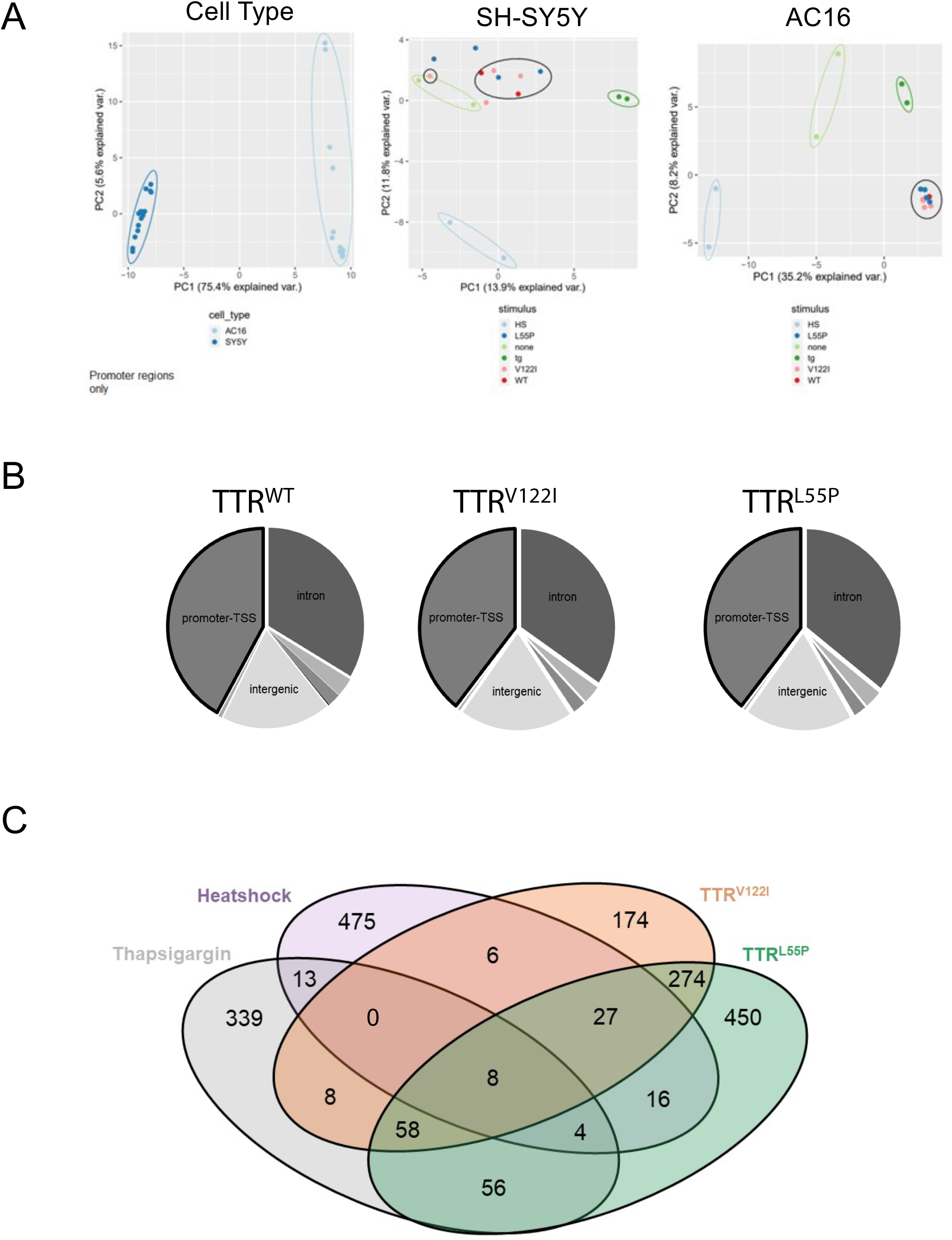
SH-SY5Y and AC16 cells exhibit changes in chromatin accessibility in response to destabilized TTR exposure and profound ER stress. **(A)** PCA plots separating all samples depicts grouping dictated by cell type (left) by changes in chromatin accessibility as compared to naïve (untreated samples). Middle and right plots represent SH-SY5Y (left) and AC16 (right) cells exposed to each stimulus. **(B)** Percentage of total DAPs in AC16 cells exposed to TTR^WT^, TTR^L55P^, or TTR^V122I^ located within promoter regions, intergenic regions, intronic regions, or other non-coding regions of the genome. **(C)** Comparison of DAPs resulting from exposure of AC16 cells to HS, thapsigargin, and destabilized TTRs.

